## Supplementary Information ver2 for "Spatiotemporal development of cuticular ridges on leaf surfaces of *Hevea brasiliensis* alters insect attachment"

**Table S1:** Age, color and surface properties of the adaxial leaves of *Hevea brasiliensis* at different ontogenetic stages

**Fig. S1:** Leaf growth versus leaf stage: The image shows the length of midrib (growth) versus leaf age for four *Hevea brasiliensis* leaves

**Fig. S2**: Schematic of the experimental set up used for insect walking experiments to measure maximum traction forces. PC – Computer, AD – Amplifier, FT – Force transducer, L – Light source, W – Molten beeswax, B – Beetle, R – Replica and H – Human hair.

**Fig. S3:** 3D reconstruction of CLSM observations of replicas of leaves at different stages using Mountains Map Premium ver. 7: (a) Stage 1 (b) Stage 2A (c) Stage 2B (d) Stage 3 (e) Stage 2B (leaf remnants attached) (f) Stage 3 (leaf remnants attached) (g) Stage 4 (h) Stage 5.

**Fig. S4:** Leaf cuticle patch attached to the epoxy replica (a) Leaf cuticle (b) Epoxy

**Fig. S5:** Correlation plot of log transformed values of mean insect traction forces versus mean aspect ratio of the ridges taken over each replicate (without data from stages S2B and S3). Pearson's test showed strong correlation of insect traction forces with mean aspect ratio of the ridges with $R = -0.91$

**Table S1: Age, color and surface properties of the adaxial leaves of *Hevea brasiliensis* at different ontogenetic stages**

| **Leaf Stage** | **Notation** | **Leaf Age** | **Leaf color** | **Surface structure** |
|---|---|---|---|---|
| Stage 1 | S1 | 13 ± 2 | Shiny brown | Smooth cells |
| Stage 2 (apical) | S2A | 15 ± 3 | Shiny brown | Smooth cells |
| Stage 2 (basal) | S2B | 15 ± 3 | Pale green | Ridges<br>(high aspect ratio) |
| Stage 3 | S3 | 16 ± 3 | Pale green | Ridges<br>(high aspect ratio) |
| Stage 4 | S4 | 21 ± 4 | Pale green | Ridges<br>(labyrinth type) |
| Stage 5 | S5 | > 60 | Dark green | Ridges<br>(labyrinth type) |

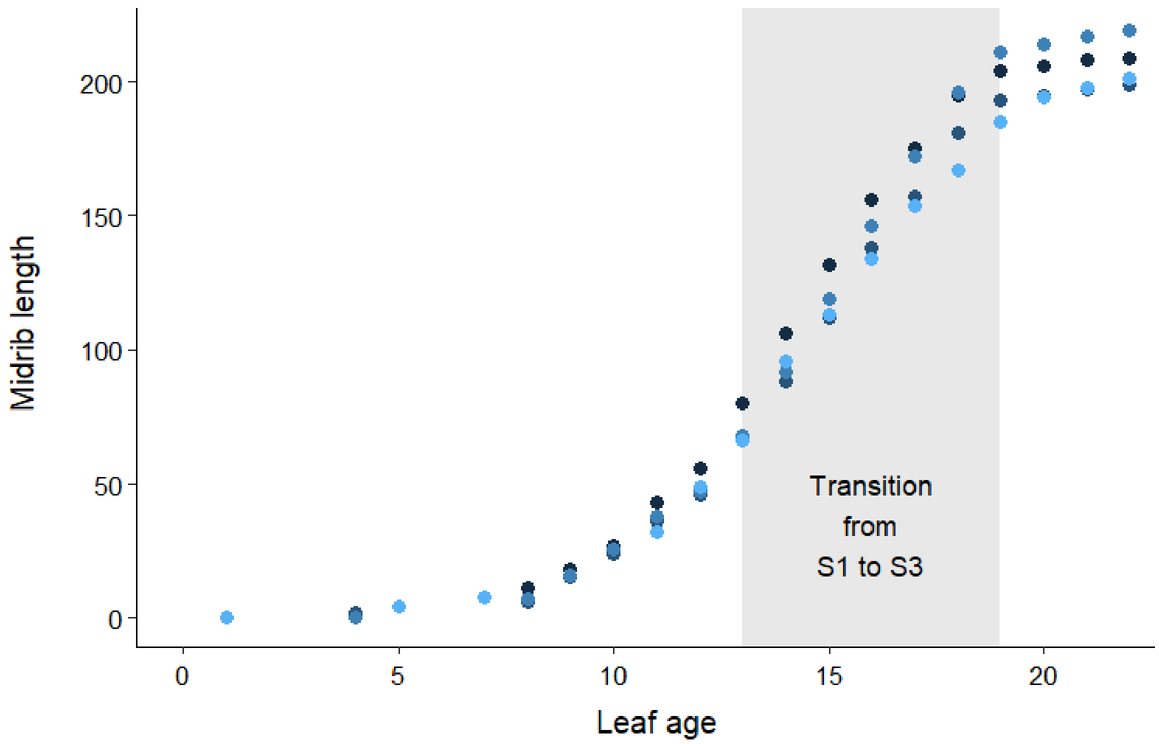


**Fig. S1: Leaf growth versus leaf age**: The image shows the length of midrib (growth) versus leaf age for four *Hevea brasiliensis* leaves

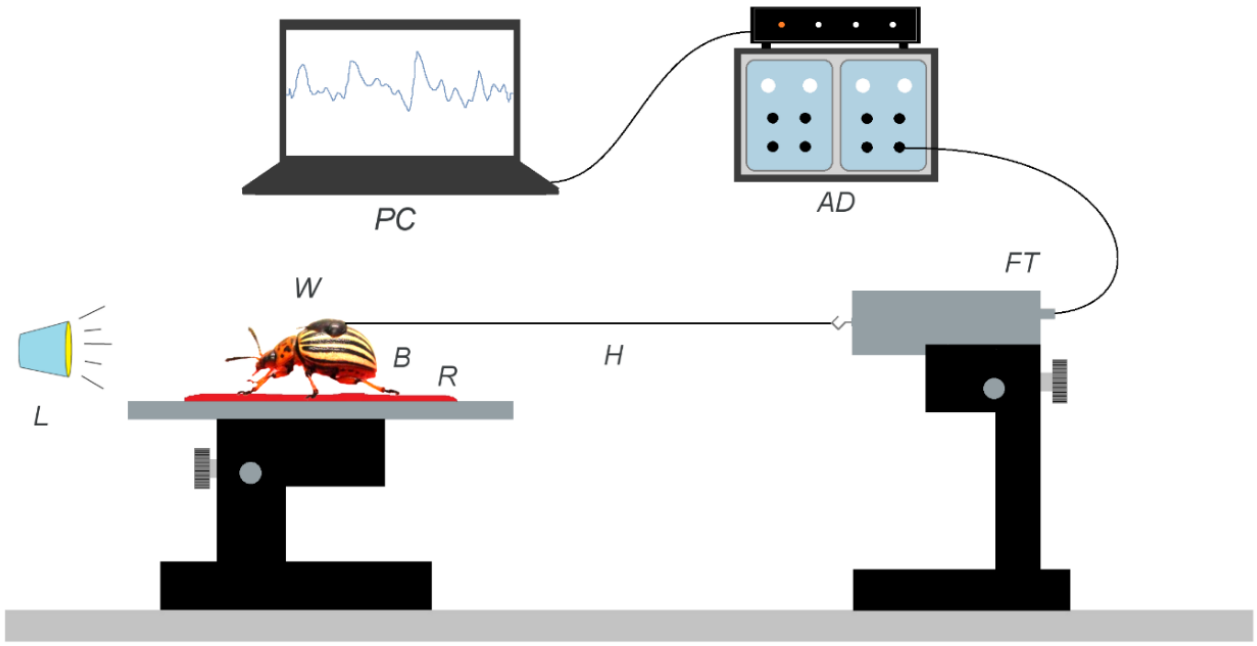


**Fig. S2**: Schematic of the experimental set up used for insect walking experiments to measure maximum traction forces. PC – Computer, AD – Amplifier, FT – Force transducer, L – Light source, W – Molten beeswax, B – Beetle, R – Replica and H – Human hair.

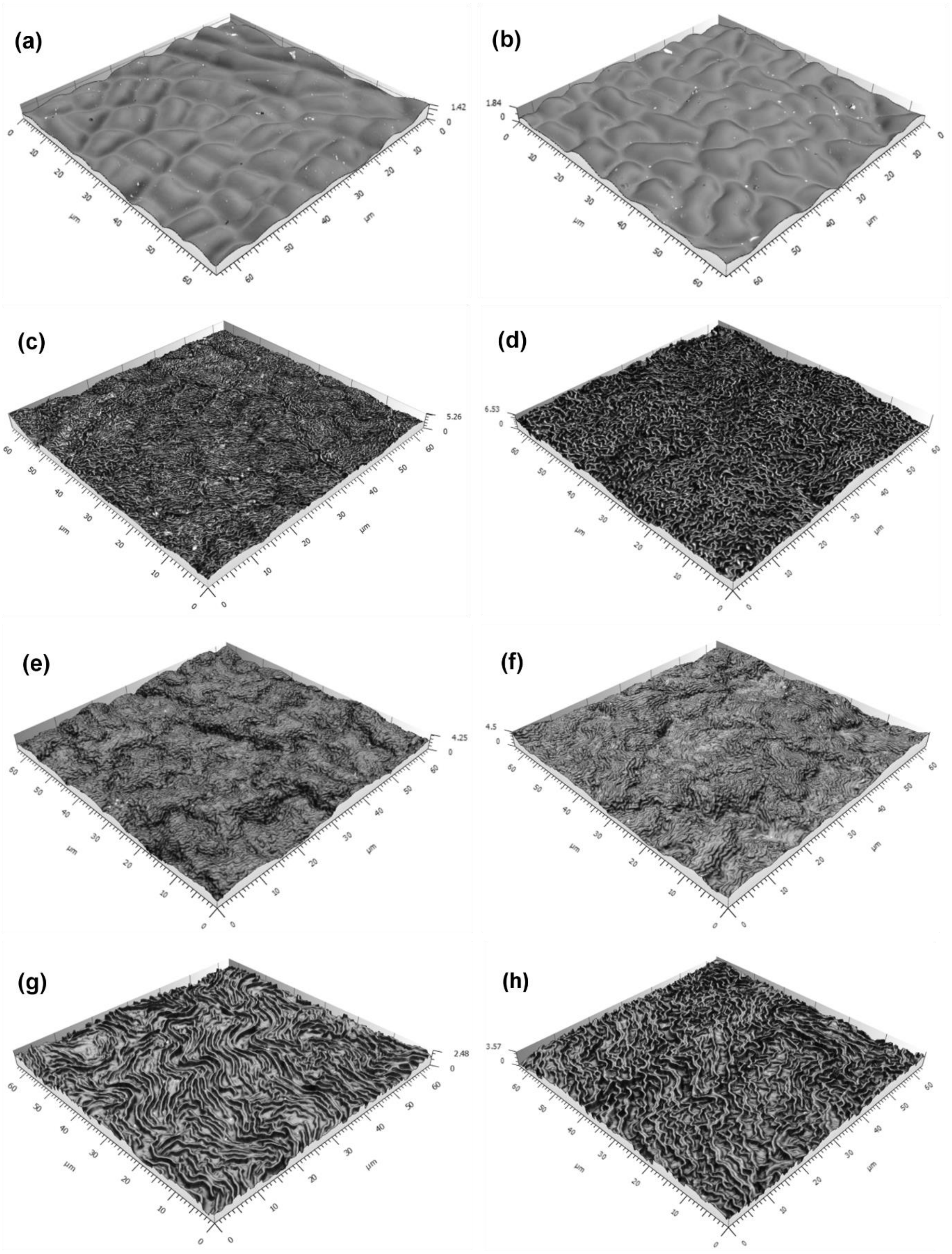

**Fig. S3: 3D reconstruction of CLSM observations of replicas of leaves at different stages using Mountains Map Premium ver. 7:** (a) Stage 1 (b) Stage 2A (c) Stage 2B (d) Stage 3 (e) Stage 2B (leaf remnants attached) (f) Stage 3 (leaf remnants attached) (g) Stage 4 (h) Stage 5.

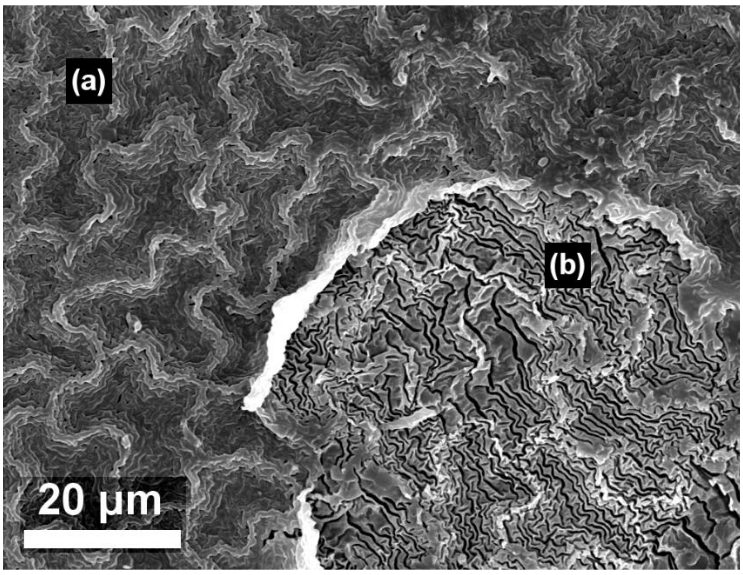


**Fig. S4:** Leaf cuticle material attached to the (negative) epoxy replica (a) Leaf cuticle (b) Epoxy

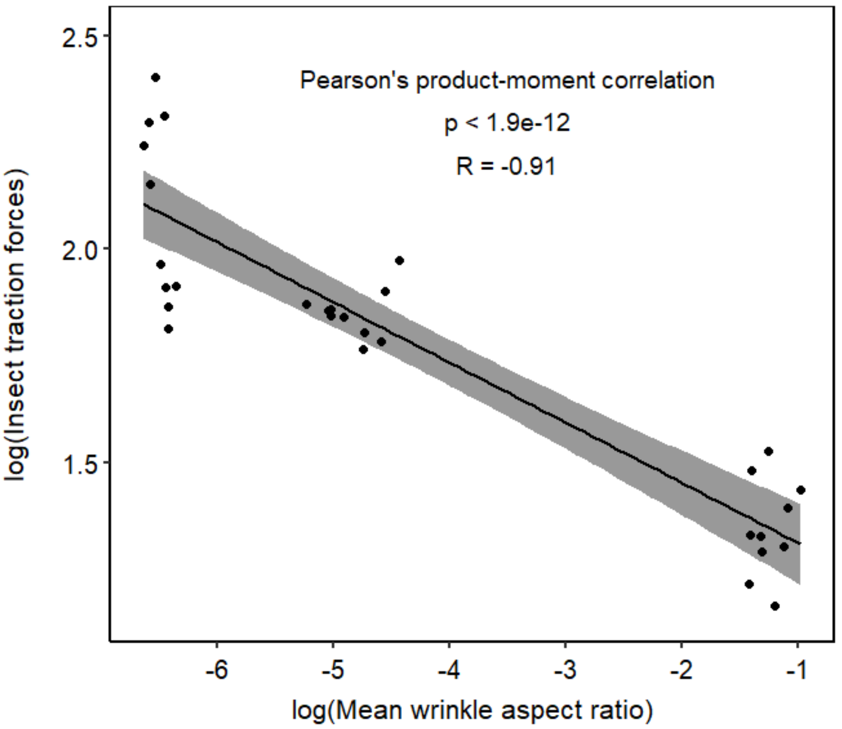


**Fig. S5:** Correlation plot of log transformed values of mean insect traction forces versus mean aspect ratio of the ridges taken over each replicate (without data from stages S2B and S3). Pearson's test showed strong correlation of insect traction forces with mean aspect ratio of the ridges with $R = -0.91$
